# Open microfluidic coculture reveals paracrine signaling from human kidney epithelial cells promotes kidney specificity of endothelial cells

**DOI:** 10.1101/2020.02.14.949776

**Authors:** Tianzi Zhang, Daniel Lih, Ryan J. Nagao, Jun Xue, Erwin Berthier, Jonathan Himmelfarb, Ying Zheng, Ashleigh B. Theberge

**Affiliations:** Department of Chemistry, University of Washington, Seattle, 98105, USA; Department of Bioengineering, University of Washington, Seattle, 98105, USA; Department of Medicine, University of Washington, Seattle, 98105, USA; Department of Kidney Research Institute, University of Washington, Seattle, 98105, USA; Department of Institute for Stem Cell and Regenerative Medicine, University of Washington, Seattle, 98105, USA; Department of Urology, University of Washington, Seattle, 98105, USA

## Abstract

Endothelial cells (ECs) from different human organs possess organ-specific characteristics that support specific tissue regeneration and organ development. EC specificity are identified by both intrinsic and extrinsic cues, among which, parenchyma and organ-specific microenvironment are critical contributors. These extrinsic cues are, however, largely lost during *ex vivo* cultures. Outstanding challenges remain to understand and re-establish EC organ-specificity for *in vitro* studies to recapitulate human organ-specific physiology. Here, we designed an open microfluidic platform to study the role of human kidney tubular epithelial cells in supporting EC specificity. The platform consists of two independent cell culture regions segregated with a half wall; culture media is added to connect the two culture regions at a desired timepoint, and signaling molecules can travel across the half wall (paracrine signaling). Specifically, we report that in the microscale coculture device, primary human kidney proximal tubular epithelial cells (HPTECs) rescued primary human kidney peritubular microvascular EC (HKMEC) monolayer integrity and fenestra formation, and HPTECs upregulated key HKMEC kidney-specific genes (*HNF1B*, *AJAP1*, *KCNJ16*) and endothelial activation genes (*VCAM1*, *MMP7*, *MMP10*) in coculture. Co-culturing with HPTECs also promoted kidney-specific genotype expression in human umbilical vein ECs (HUVECs), and human pluripotent stem cell-derived ECs (hPSC-ECs). In comparison to the culture in HPTEC conditioned media, co-culture of ECs with HPTECs showed increased upregulation of kidney specific genes, suggesting potential bidirectional paracrine signaling. Importantly, our device is compatible with standard pipettes, incubators, and imaging readouts, and could also be easily adapted to study cell signaling between other rare or sensitive cells.

## Introduction

During human development, ECs acquire tissue-specific properties based on cues from the local microenvironment while organs undergo vascularization; *in vitro* cell culture systems often lack microenvironmental cues, making it challenging to maintain endothelial cell identity (20). Therefore, a better understanding of how to preserve endothelial cell morphologic phenotypes and gene expression profiles in *ex vivo* environments is important to building more physiologically relevant endothelial cell-based disease models for pathological research. Specifically, the human kidney is a highly vascularized organ, and the peritubular capillaries play an important role in maintaining normal renal function of selective reabsorption and secretion, in addition to providing oxygen and nutrients to the tubules and surrounding cells (1, 27, 33). The loss of renal microvasculature integrity has been recognized as a classic finding contributing to progressive renal diseases, including tissue ischemia, tubular dysfunction, inflammation, and fibrosis (27, 32, 36). Reabsorption occurs through both passive and active transport, from the proximal tubular epithelial cells lining the tubule, across the extracellular matrix (ECM) separating the tubules, through the ECs and into the adjacent peritubular capillaries (9).

The interplay between human kidney proximal tubule epithelial cells (HPTECs) and human kidney peritubular microvascular ECs (HKMECs) is a complex process largely regulated by soluble factors in the tubulo-interstitium (27, 34, 38). The understanding of human kidney microvasculature injury and
regeneration mechanisms has been of great interest in the past several decades, and a group of growth factors have been identified that mediate the intercellular interaction (52, 53, 55). Importantly, vascular endothelial growth factor (VEGF) is an angiogenic and vascular permeability factor, which is critical to the survival and proliferation for ECs (23, 28). However, due to the challenge of isolating primary HKMECs from fresh tissues and the unavailability of an established HKMEC cell line, much of our understanding has come from cell culture models using non-renal human cell lines, such as human umbilical vein ECs (HUVECs), or animal cells. For example, Kim et al (28) showed that human kidney epithelial cells generate VEGF which promoted branching angiogenesis of HUVECs *in vitro*; Zhao et al. (62) discovered that mouse renal proximal tubular epithelial cells helped mouse renal peritubular ECs maintain phenotypes in coculture. More recently, our group (33) successfully isolated and purified HKMECs and discovered that the significant difference between HKMECs and HUVECs in morphology, phenotype and transcriptional profiling. Further, we (37) reported that ECs from different human organs exhibit organ-specific gene expression profiles, which correlate with specific cell functions, including metabolic rate, angiogenic potential, and barrier properties. However, compared to freshly isolated cells, most of organotypic gene expression is lost during *in vitro* expansion, which highlighted the importance of both employing specific endothelial cell types and identifying ways to enhance organotypic properties during *in vitro* culture.

We sought to test whether soluble factors from HPTECs could be used to maintain kidney-specific gene expression and morphology in HKMECs cultured *in vitro* using a microfluidic coculture platform, and further to understand if non-parenchymal tissues (i.e., non-kidney ECs) are plastic to develop organ-specificity through paracrine signaling with parenchymal cells (i.e., kidney epithelial cells). Emerging as an alternative tool for coculture studies, microfluidic platforms reduce cell and reagent consumption and offer better control over the configuration of the cell culture regions. Despite the rapid development of microfluidic technologies, many microfluidic platforms require handling expertise and equipment such as pumps and valves, creating significant obstacles for general biological laboratories to adopt microfluidic cell culture devices (8, 48, 61). In this study, our goal was to create a user-friendly microfluidic device not only suitable for our coculture objectives, but also translatable to other cell culture schemes and easily adoptable by general biology researchers.

Here, we present an open microscale coculture platform that contains two cell culture chambers segregated by a polystyrene half wall. Paracrine signaling is initiated by connecting the two chambers with additional cell culture media, which overflows the half wall. We separately cultured HPTECs with three different endothelial cell types in the device and evaluated key endothelial cell organ-specific features under differential culture conditions. Our findings suggest that coculture with HPTECs supports maintenance of key organ-specific genes in primary human kidney ECs (HKMECs) and enhances kidney-specific gene expression in non-kidney-derived cells (human umbilical vein ECs (HUVECs), and human pluripotent stem cell-derived ECs (hPSC-ECs) over three days in culture. This work also serves as a proof of concept to demonstrate that the presented device could become a simple and efficient platform to study paracrine signaling effects across different cell types.

## Results and discussion

### Open microfluidic device design and workflow

Open microfluidic systems are characterized by the introduction of an air-liquid interface, where liquid flows in microscale channels devoid of at least one side-wall (2, 5, 6, 13). Unlike typical microfluidic devices with enclosed culture chambers, open devices offer increased pipette accessibility at any point along the culture chamber and simpler fabrication processes, such as straightforward micro-milling and injection molding, allowing open microfluidic devices to become efficient tools with versatility and transferability across different research disciplines (11, 18, 30, 31, 40, 49, 58).

In the present study, we develop a user-friendly platform to maintain and evaluate kidney endothelial cell organ-specificity under various microenvironments. Our device consists of two cell culture regions, a center chamber and a side chamber, segregated by a half wall (Fig. 1A, Fig. S1A). The width and height of the side chamber are designed to allow filling by a single pipette-dispensing step based on the concept of spontaneous capillary flow in open microchannels (see SI for equation describing the conditions for capillary flow and illustrations in Fig. S1, Video S1). Two cell types are selectively pipetted into the center (ECs) and the side (HPTECs) chamber, respectively; after adherence, the cells are placed into paracrine signaling contact by filling cell culture medium over the half wall (Fig. 1B). The notched features for each chamber are designed to accommodate a pipette tip for ease of use when pipetting media. Fig. 1C shows a device with 5 by 4 array of the wells. The replicating wells allow for testing multiple culture conditions in a single experiment. Additionally, the dimensions of the device plate are designed to fit into a standard petri dish (88 mm in diameter, shown in Fig. 1C) to maintain sterility and control evaporation during incubation. The device plate dimensions (60×50 mm) are also compatible with universal microscope mounting frames for convenient endpoint imaging.

**Figure 1:**
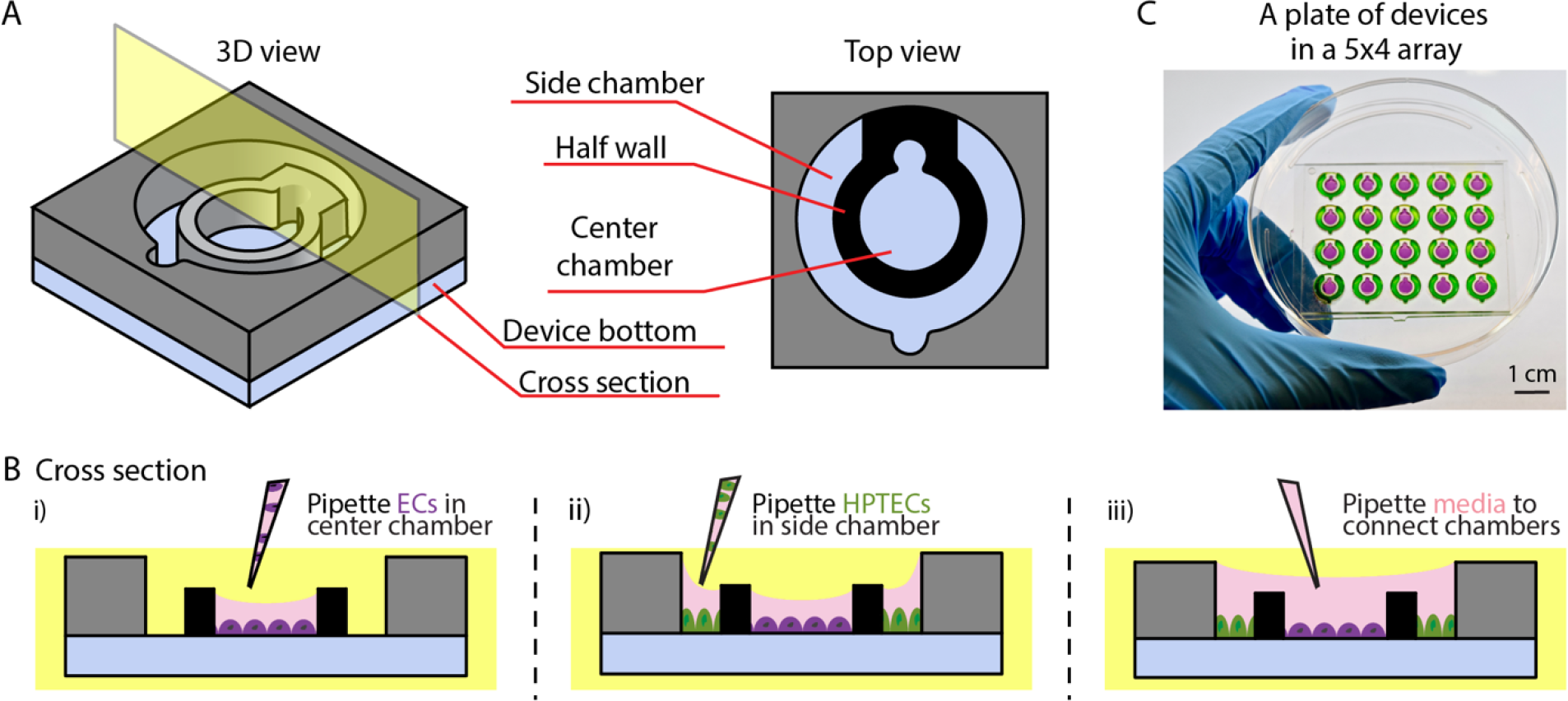
Coculture device design and operation. (A) 3D and top view of device design. (B) Cross section view of device operation: different cell types (endothelial cells (ECs) and HPTECs) are selectively seeded into the separated cell culture chambers (i-ii) and placed into paracrine signaling contact by addition of shared media on top (iii). (C) A photograph of a plate of devices in 5×4 array fitted inside a petri dish. Center and side chambers are loaded with purple and green dye, respectively, to visualize chamber segregation. The device loading process is shown in Video S1.

Importantly, the coculture device is fabricated with polystyrene, which is the standard material for cell culture due to its biocompatibility and good optical properties for medium resolution imaging, which is used for standard kidney endothelial cell immunocytochemistry and morphology studies (17). In contrast to microfluidic platforms made of polydimethylsiloxane (PDMS), the device used in this work mitigates problems with oligomer leaching and adsorption of small molecules (such as potential drugs or toxicants that this device could be used to screen in future studies) (7, 25, 45, 54). The ECs (HKMECs, HUVECs, and hPSC-ECs) and HPTEC coculture model presented in this study demonstrates that the coculture device, and our culture protocols to avoid evaporation and ensure nutrient availability in microscale culture, are compatible with cells from different sources. Additionally, to demonstrate the versatility of the device for culturing different cells in the side chamber, we cultured freshly isolated human pericytes in the side chamber and HKMECs in the middle chamber for 72 hours; this is a biologically relevant coculture model because kidney pericytes play a profound role in inflammation and the renal pericyte-endothelial cross-talk contributes to fibrogenesis during kidney disease progression (29).The results showed that incorporation of pericytes increased HKMEC density in the device (Fig. S2).

### HKMEC and HPTEC coculture in an open microscale device preserves endothelial morphology

A central goal of our study is to understand to what extent EC specificity can be modified by parenchymal cells in vitro, and potentially to develop a method to maintain kidney specificity of HKMECs *in vitro.* The loss of microvascular endothelium in renal disease progression is directly related to altered local secretion of VEGF *in vivo*, which is mainly produced by proximal tubular epithelium (21, 46, 57). We utilized the open microscale coculture devices to examine the morphology of HKMECs in four different cultural conditions (Fig. 2A). Briefly, HKMECs were cultured in the center chambers of the device alone or with HPTECs cultured in the side chambers, both in media with and without exogeneous VEGF. We used immunocytochemistry to qualitatively compare the expression of vascular endothelial cadherin (VECad), an endothelial integrity marker that is an essential component of endothelial intercellular adherens junctions and critical to endothelium integrity and barrier function (12), and the expression of plasmalemma vesicle associated protein (PV1), a marker critical to the formation of stomatal (in the peri-nuclear cytosol) and fenestral (in the periphery of EC membranes) diaphragms on endothelial cell membranes (50).

**Figure 2:**
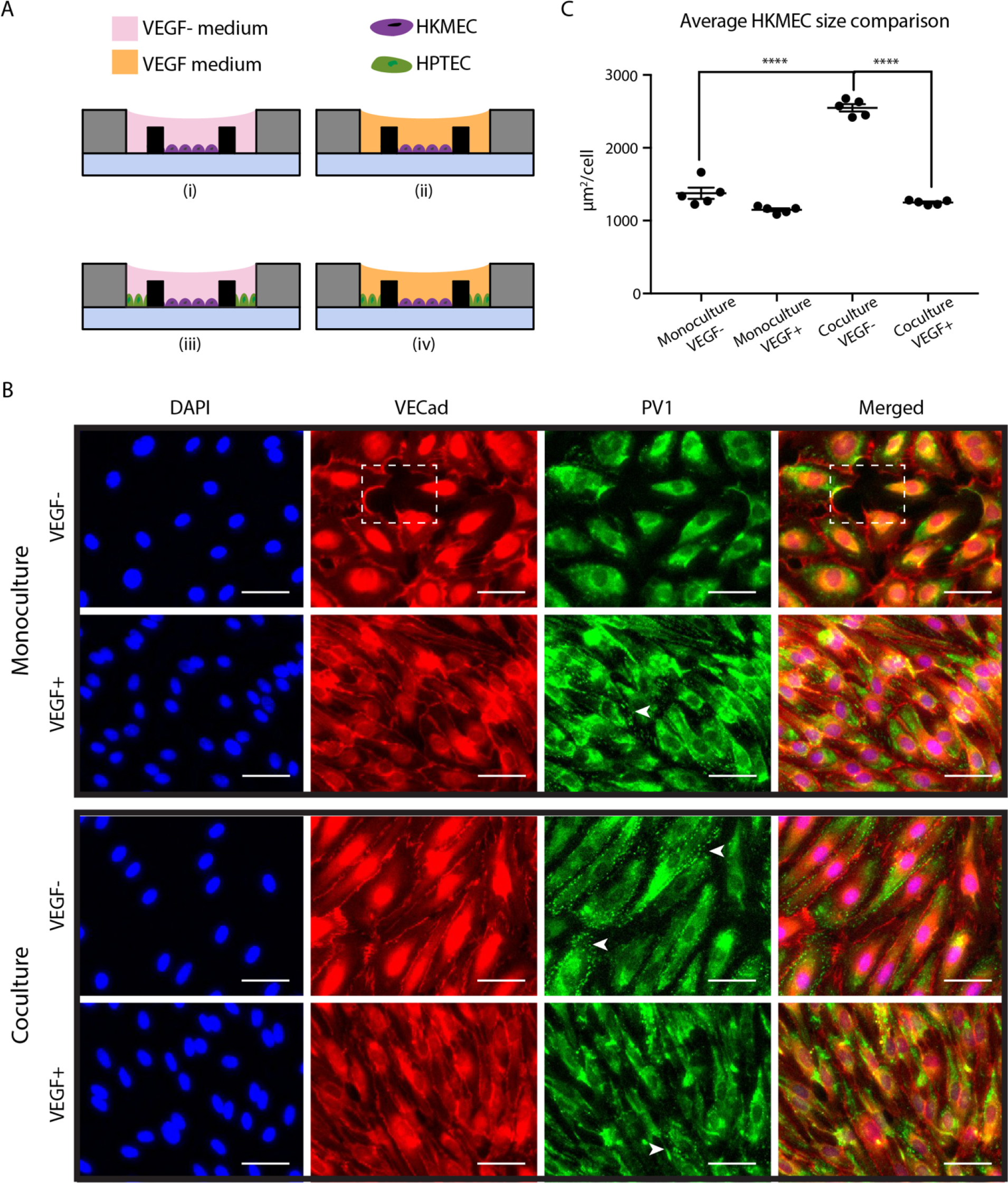
Coculture with HPTECs preserves HKMECs morphology in coculture. (A) Device cross section views showing the four culture conditions of HKMECs and HPTECs: monoculture of HKMECs in media without (i) or with (ii) exogeneous VEGF; segregated coculture of HKMECs and HPTECs in media without (iii) or with (iv) exogeneous VEGF. (B) Immunofluorescence images of HKMECs under differential culture conditions. Blue: nuclear stain (DAPI). Red: vascular endothelial cadherin (VECad); dotted box indicates a region without cells. Green: plasmalemma vesicle associated protein (PV1); arrows indicate example regions with clear PV1 structures (punctate staining). Scale bars: 50 μm. Images shown are from donor C and are representative of experiments with cells from 3 different human donors each with 5 device replicates per experiment. (C) Quantification of HKMEC size under differential culture conditions. Error bars represent the SEM of five device replicates for a single human donor (donor C) in a single experiment. Data sets were analyzed using a one-way ANOVA test; p-values are indicated for Tukey’s multiple comparisons tests between “Coculture, VEGF-” and “Monoculture, VEGF-” conditions; and “Coculture, VEGF-” and “Coculture, VEGF+” conditions. (\*\*\*\**p* ≤ 0.0001)

As shown in Fig. 2B, without exogenous VEGF, HKMECs cultured alone (i.e., “Monoculture, VEGF-”) lost consistent expression of VECad between adjacent cells, and gaps between the cells were observed (Figure 2B, box), indicating disrupted EC barrier. Additionally, in the “Monoculture, VEGF-” condition, majority of PV1 expression is perinuclear, suggesting limited fenestrae formation. Addition of exogenous VEGF in monoculture enabled a contiguous monolayer without gaps (as visualized in the VECad staining) and partially rescued PV1 expression. With the presence of HPTECs in VEGF-free media (i.e., “Coculture, VEGF-”), HKMECs maintained integrated cell-cell contact (VECad staining) and showed clusters of clear PV1 expression (Figure 2B, arrows, punctate staining), indicating that soluble factors from the neighboring but physically separated HPTECs helped maintain important endothelial cell function without exogenous VEGF. In addition, HKMECs in the “Coculture, VEGF-” condition had larger cell size when compared to the “Monoculture, VEGF-”, and “Coculture, VEGF+” conditions (*p*<0.0001, Fig. 2C). This suggests that HPTECs co-culture promoted HKMECs quiescence and maturation (35). Adding VEGF alone did not have the same influence as the co-culture condition, suggesting that PTECs affect HKMECs via different or additional small molecule signaling than VEGF alone. The similar trend was observed across donors, and additional data from donor A is provided in Fig S3. Taken together, our results show the utility of the coculture device as an *in vitro* platform able to recapitulate key kidney endothelial protein expression and morphology sustained by soluble factor crosstalk with kidney epithelial cells.

### HPTECs support expression of key HKMEC organ-specific and activation genes in coculture

Previous studies observed that during organ development, ECs from different organs are heterogeneous in both morphology and gene expression patterns and participate in crosstalk with surrounding microenvironments to form the organ-specific vasculature (14, 39). Multiple kidney-specific endothelial genes are highly expressed in freshly isolated primary kidney endothelial tissues yet become significantly downregulated in expanded culture (37). We wanted to determine if coculture with HPTECs could help rescue this loss in parenchymal gene expression by soluble factor signaling. With the same setup shown in Figure 2A, we cultured HKMECs under the four different conditions (Figure 2A) and collected RNA for quantitative gene expression analysis (RT-qPCR) of selected genes.

We chose to focus on a panel of genes that were previously identified as representative molecular markers for HKMECs (37). The qPCR data in Fig. 3 indicate that compared to HKMECs in monoculture VEGF-free media (red bars), HKMECs in coculture VEGF-free media (green bars) exhibited upregulation of selective kidney-specific genes (top panel), including a 3 to 21-fold increase in Hepatocyte Nuclear Factor 1 Homeobox B (*HNF1B*), a transcription factor important to nephron development; a 7 to 28-fold in Adherens Junctions Associated Protein 1 (*AJAP1*), an adhesion junctional protein precursor crucial to endothelium barrier function; and a 2 to 32-fold in Potassium Voltage-Gated Channel Subfamily J Member 16 (*KCNJ16*), a gene encoding potassium channel protein essential to fluid regulation and pH balance. Key endothelial activation genes (bottom panel) followed a similar trend of upregulation in the coculture (green bars) versus monoculture (red bars) condition, including a 33 to 95-fold increase in *MMP7* and a 2 to 13-fold increase in *MMP10*, two matrix metallopeptidases, and a 6 to 526-fold increase in Vascular Cell Adhesion Molecule 1 (*VCAM1*), which regulates leucocyte adhesion to vascular wall (15, 19, 26, 42, 51). Additionally, HKMECs from donor C (square symbol) showed 5-fold upregulation of kidney specific gene Contactin 4 (*CNTN4*), which is linked to normal kidney development, and a 6-fold upregulation in endothelial activation gene Angiopoietin 2 (*ANG2*), which regulates angiogenesis and vascular inflammation in coculture versus monoculture (24). Interestingly, HKMECs cocultured in VEGF-supplemented media (blue bars) showed less upregulation on average of the aforementioned genes except for *ANG2*, as compared to HKMECs cocultured in VEGF-free media (green bars). These data suggest that in coculture, HPTECs secrete not only VEGF but also other soluble factors supporting HKMECs to maintain key kidney endothelial features via paracrine signaling (22, 43, 56).

**Figure 3:**
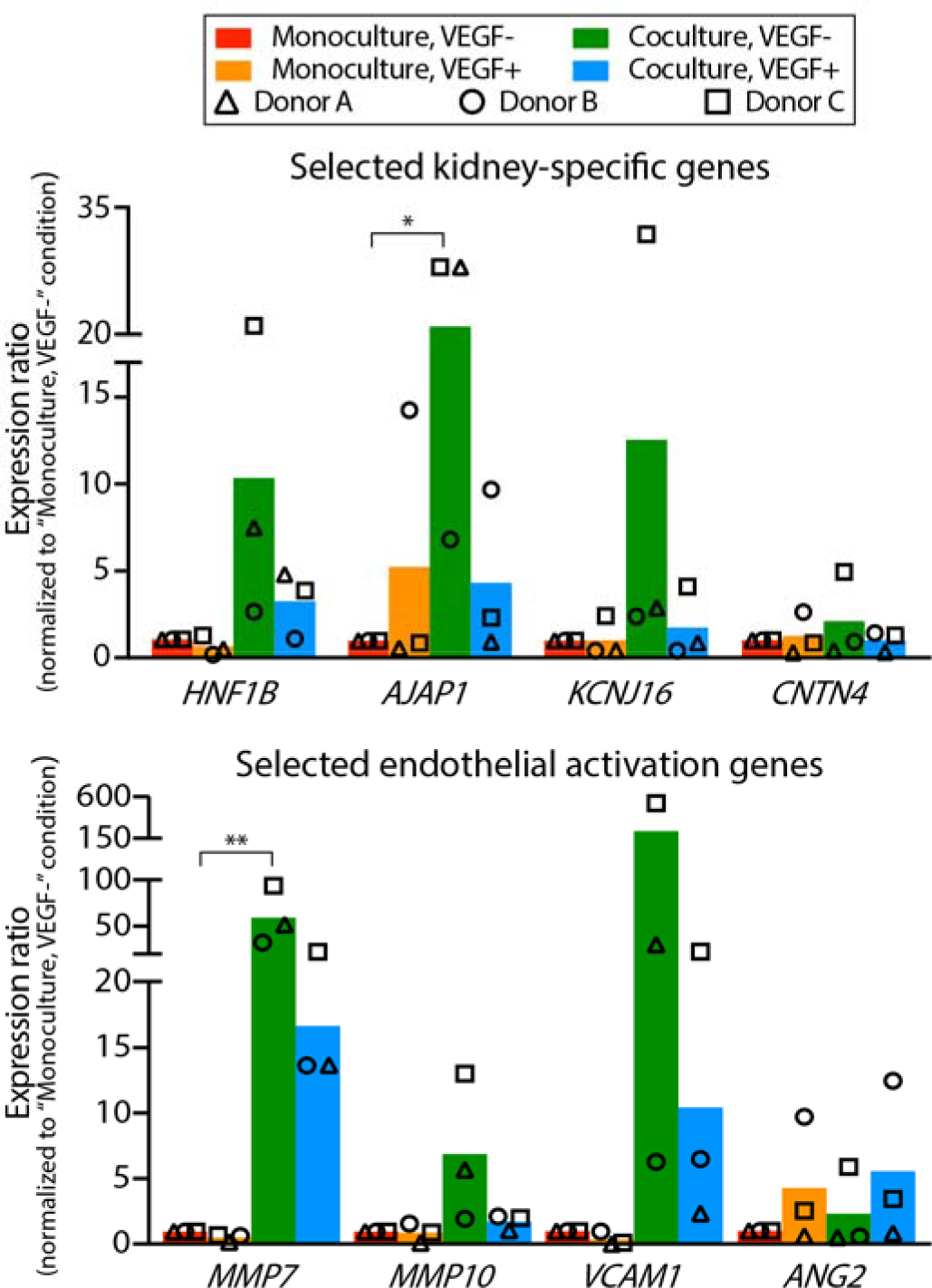
HPTECs support expression of key HKMEC organospecific and activation genes in coculture. RT-qPCR of selected HKMEC-specific genes (top) and activation genes (bottom) show upregulation in “Coculture, VEGF-” condition (green) compared to “Monoculture, VEGF-” condition (red). Each plotted point represents data from an independent human donor (pooled from 5 replicate microculture devices per donor, average of 3 RT-qPCR technical replicates), with each normalized to the “Monoculture, VEGF-” condition (which is set to 1). Each colored bar represents the average ratio of the three donors. Statistical comparisons (as described in the methods section) are shown for genes showing significant differences between “Coculture, VEGF-” and “Monoculture, VEGF-” as described in the methods section (\**p* ≤ 0.05, \*\**p* ≤ 0.01).

In contrast to the genes discussed in Fig. 3, a subset of genes including *PV1*, *VECad*, Nitric Oxide Synthase 3 (*NOS3*, an endothelial proliferation gene), and VEGF Receptor 2 (*VEGFR2*) were upregulated when cultured in “VEGF+” media compared to “VEGF-” media, for both monoculture (orange vs. red bars) and coculture (blue vs. green bars) conditions (Fig. S4). The result suggested the critical role of VEGF in HKMEC proliferation and certain aspects of kidney function development.

Figure 3 and S4 also demonstrate variance across different human donors, which highlights the importance of studying human cell signaling pathways with patient specific cells to understand patient-dependent disease mechanisms. Due to the variation across human donors, only *AJAP1* and *MMP7* show statistical significance between “Monoculture, VEGF-” condition (red bar) and “Coculture, VEGF-” condition (green bar); however, from a biological perspective, it is meaningful that all three human donors show at least 2-fold higher expression in the latter than the former condition for *HNF1B*, *AJAP1*, *KCNJ16*, *VCAM1*, *MMP7*, and *MMP10*. The response among donors for *CNTN4* and *ANG2* was mixed, with only donor C (square symbol) showing upregulated expression. Moreover, donor C showed the most pronounced upregulation in “Coculture, VEGF-” condition than the other two donors across all genes of interest in Fig 3.

### HPTECs induce key kidney-specific markers in other types of ECs

As shown in the preceding figures, HKMECs display plasticity in the coculture device and demonstrate that paracrine signaling with HPTECs led to enhanced parenchymal properties. Further, we wanted to investigate if parenchymal cells (i.e., HPTECs) have inherent potential to tune human non-kidney ECs through paracrine signaling.

Human umbilical vein ECs (HUVECs) have been widely used as a microvascular endothelial cell surrogate in traditional kidney research (19, 26, 28, 52, 53). We determined whether coculture with HPTECs could induce parenchymal changes (i.e., kidney-specific expression) in HUVECs through paracrine signaling. We placed HUVECs in the center chamber of the coculture device with or without HPTECs seeded in the side chamber. After three days of incubation, we used immunocytochemistry to evaluate expression of the platelet-endothelial cell adhesion molecule (PECAM1, also known as CD31), blood glycoprotein von Willebrand Factor (vWF), and actin cytoskeleton molecule (phalloidin staining) in differential culture conditions (16, 47). Fig. 4A shows that HUVECs in monoculture had robust PECAM1 localization at intercellular junctions, whereas in coculture, very low levels of PECAM1 resided at the intercellular junctions, suggesting junctional instability and cell activation. In addition, HUVECs in coculture showed less vWF in cell cytosol than in monoculture, and more aligned phalloidin structure, suggesting that HPTECs perturbed endothelial quiescence and led to endothelial activation (4). Together, the data suggest that HPTEC perturbed HUVEC quiescence in coculture, and HPTECs secreted soluble factors that may stimulate endothelial activation in HUVECs.

**Figure 4.**
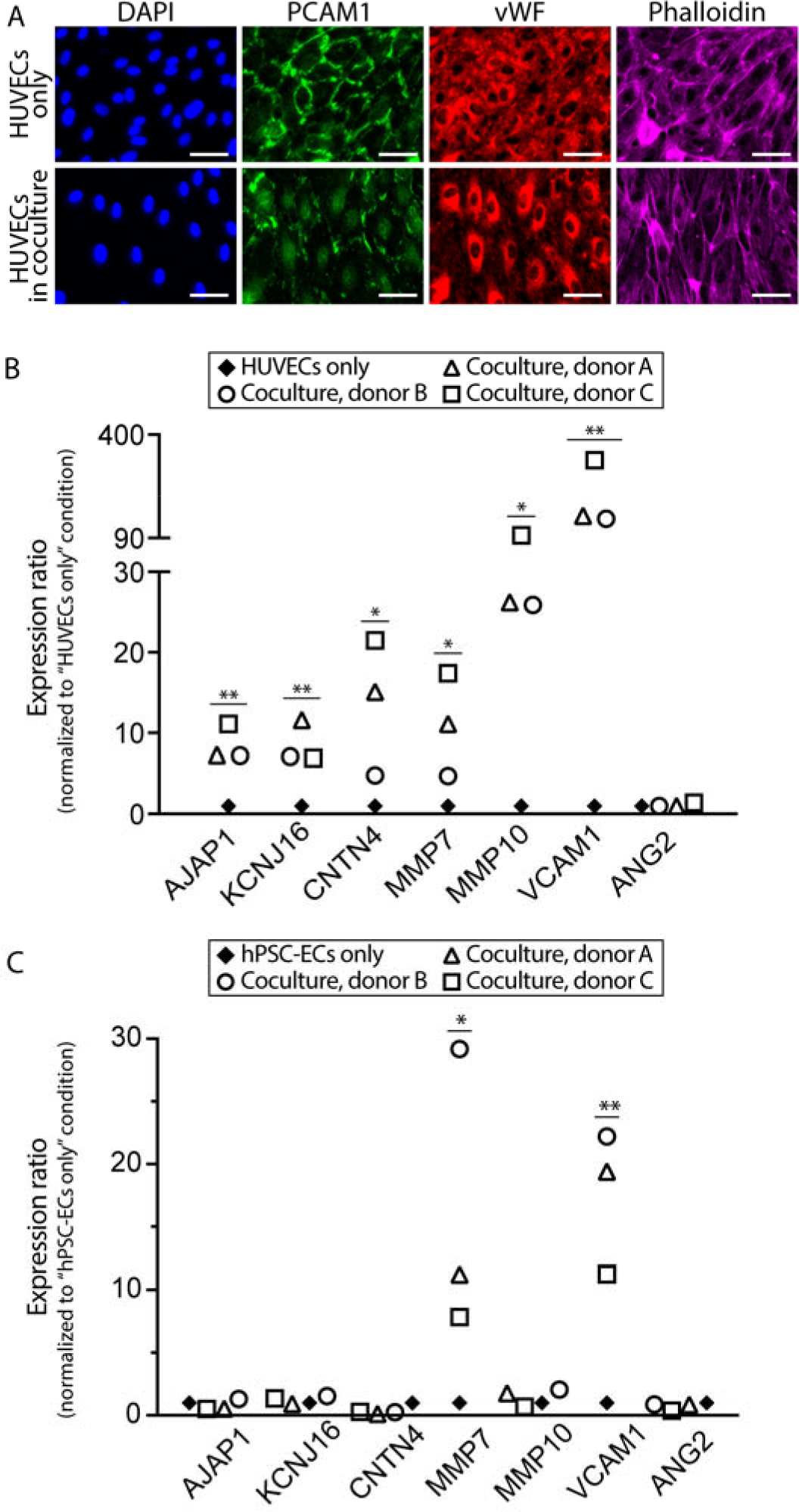
HPTECs activate other types of endothelial cells in coculture. (A) Immunofluorescence images of HUVECs in monoculture and coculture condition with HPTECs. Scale bars: 50 μm. (B) Selected kidney-specific markers and endothelial activation markers in HUVECs cocultured with HPTECs from three different donors. (C) Selected kidney-specific markers and endothelial activation markers in hPSC-ECs cocultured with HPTECs. The RT-qPCR samples were pooled from 5 replicate microculture devices per donor, average of 3 RT-qPCR technical replicates. Gene expression in both (B) and (C) is normalized to monoculture, which is set to 1. Statistical comparisons (as described in the methods section) are shown for genes showing significant differences between coculture and monoculture (\**p* ≤ 0.05, \*\**p* ≤ 0.01).

We also compared the expression of the same panel of kidney-specific and activation genes in HUVECs cultured alone or with HPTECs. Fig. 4B suggests that similar to HKMECs in coculture with HPTECs, HUVECs expressed higher levels of the kidney organ-specific genes, including *AJAP1* (7 to 11-fold), *KCNJ16* (7 to 12-fold), and *CNTN4* (5 to 22-fold), and the endothelial activation genes, including *MMP7* (5 to 17-fold), *MMP10* (26 to 99-fold), and *VACM1* (148 to 323-fold) in coculture. However, none of the donor cells led to upregulation of *ANG2*. The expression of *HNF1B*, a kidney-specific gene was also detected in the HUVECs after coculture, whereas no detectable expression in the HUVEC alone condition (data not shown). Consequently, HPTECs showed potential to induce parenchymal characteristics in HUVECs through paracrine signaling.

Generating ECs *de novo* by using human pluripotent stem cells (hPSC) has emerged as a tool in the past decade to study vascular pathogenesis and construct tissues for drug screening and other therapeutic applications (32, 41). Here, we explored whether hPSC-derived ECs (hPSC-ECs) can develop kidney specific features when co-cultured with HPTECs. Briefly, we differentiated hPSCs to ECs following the protocol adapted from Palpant et al. (41), and cultured hPSC-ECs in our microscale device in center well alone (monoculture) or co-cultured with HPTECs. We show that compared to in monoculture, hPSC-ECs in coculture exhibited upregulation of *MMP7*, and *VACM1*, whereas no obvious changes of *AJAP1*, *KCNJ16*, *CNTN4*, *MMP10*, and *ANG2* across the three human donors (Fig. 4C). Similar to HUVECs, the expression of *HNF1B* was not detected in the hPSC-EC monoculture sample, but at a detectable level in the coculture samples (data not shown). Our findings suggest that parenchymal tissue coculture has the potential to tune human generic cells to develop organ-specific physiological features.

It should be noted that cross contamination of the cell types could be a potential issue during the media changing step, as unattached cells or debris could be flushed into the other cell culture chamber when fresh media is pipetted directly from the top of the device; if this occurred, the purity of the cell lysate sample used for qPCR analysis would be affected. In this study, since the immunofluorescence images of the ECs (Fig. 2) did not show any presence of the HPTECs, we are confident about the purity of the endothelial cell lysate collected from the center chamber. Further, we have explored this co-culture effect via conditioned media cultures as below, to further support the changes of ECs by HPTECs.

### HPTEC coculture and conditioned media culture lead to different effects

Historically, transferring conditioned media from one cell type to another has been a common approach to recapitulating cell-cell interactions via soluble signaling mediators. Although conditioned media experiments have facilitated the understanding of cellular signaling mechanisms and the discovery of biomarkers and therapeutic targets in diseases, they are inadequate to capture the effects of fast-decaying factors or dynamic bidirectional signaling due to the temporal segregation of the cell populations (8, 59).

Using the segregated coculture device, we were able to compare the difference between HKMECs in shared media coculture with HPTECs and HKMECs in HPTEC conditioned media culture. To model conditioned media culture, HPTECs were plated in the side chamber of the device overnight, and conditioned media was collected to feed HKMECs seeded in the center chamber of a separate device. The HPTEC conditioned media was collected every 24 hours, and the freshly collected medium was used to replace HKMEC culture medium. HKMECs were collected for RT-qPCR analysis after three days of culture with HPTEC conditioned medium.

As shown in Fig. 5, HKMECs in conditioned media culture (red) showed similar trend with noticeable upregulation of kidney specific genes, particularly *KCNJ16*, *MMP7*, *VCAM1*, whereas the changes of *HNF1B*, *AJAP1*, and *MMP10* are evidently less pronounced in conditioned media culture (red) than in coculture (green). This data suggests that the signaling between HKMECs and HPTECs involves both unidirectional signaling (in conditioned media) and bidirectional signaling (in shared media) pathways. It is also possible that some of the key signaling factors are short-lived and are therefore lost in the conditioned media experiments while still present in the coculture experiments. Our microscale device enables bidirectional signaling molecule interactions in real time, which may result in more biomimetic effects than unidirectional signaling in conditioned media studies.

**Figure 5.**
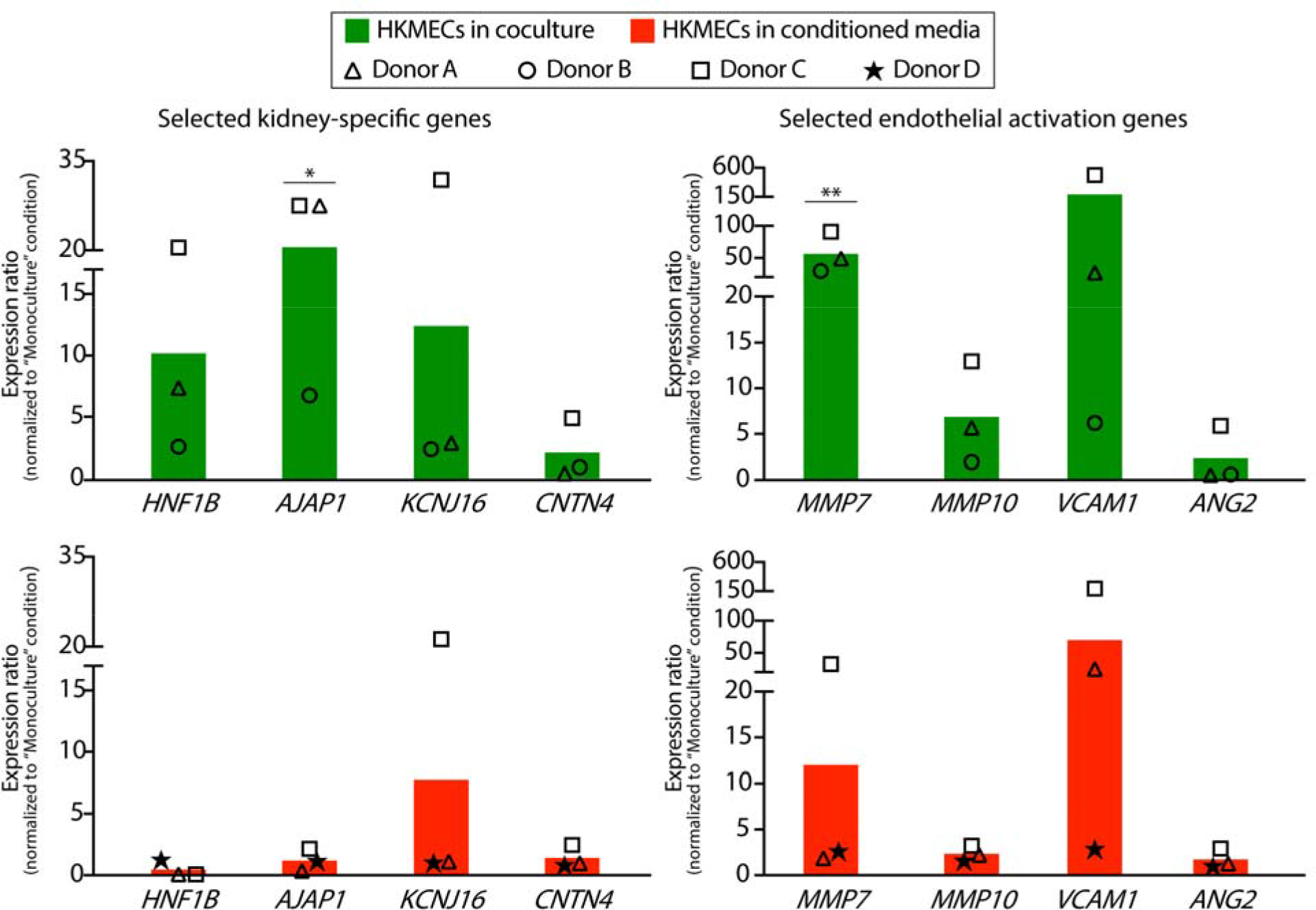
Selected gene expression profiles for HKMECs in coculture and conditioned media culture. HKMECs showed higher upregulation of kidney-specific (left) and endothelial activation (right) genes in coculture with HPTECs (green) compared to in HPTEC conditioned media (red). Each plotted point represents data from an independent human donor (pooled from 5 replicate microculture devices per donor, average of 3 RT-qPCR technical replicates). Each colored bar represents the average of three human donors. Gene expression is normalized to monoculture, which is set to 1.

## Conclusion

Here, we present an open microscale coculture device that provides spatial segregation for different cell types and supports paracrine signaling in the shared microenvironment. With relatively small cell samples and regular pipetting actions (no sophisticated external pumps or valves), a coculture configuration with operation efficiency can be achieved. Using the microscale device, we cocultured HPTECs with ECs from different sources. We observed that signaling molecules secreted by HPTECs enhance some level of kidney specific characteristics in generic human ECs (HUVECs and hPSC-ECs) and led to endothelial activation, indicating a supporting role of HPTECs to the functional development of ECs. Moreover, we observed higher upregulation in certain kidney specific genes and endothelial activation genes when HKMECs were cultured with HPTECs in shared media (coculture) than in conditioned media, suggesting a bidirectional signaling mechanism in the system. In future work, we envision performing proteomic analysis of the conditioned media collected from HPTEC monoculture and HPTEC-HKMEC coculture, which will provide more specific insight into the mediators of paracrine signaling in the system.

Due to the functional differences among ECs from different organs, it is important to use organ-specific ECs to understand specific vascular injury and regeneration mechanisms. For the study of the interaction between human kidney proximal tubule epithelium and peritubular microvascular endothelium, this is the first time that primary human kidney cell types are cocultured in shared media, more precisely resembling the *in vivo* microenvironment. Our findings suggest parenchymal cells have the capability to tune generic human cells *in vitro*. Our culture system is simple to use, and we have translated it to biological collaborators in other application areas who are able to operate the device in their own laboratories without assistance. In future work, we envision that our microscale device will be batch fabricated by injection molding (18, 30, 31) and easily adapted to tune parenchymal characteristics in *in vitro* cell culture studies, and construct patient-specific tissue models for pathological research and therapeutic applications.

## Materials and methods

### Open microfluidic coculture device fabrication

Devices were fabricated using a Series 3 PCNC Mill (Tormach, Waunakee, WI) or a DATRONneo (Datron Dynamics, Germany). The devices were designed with Solidworks 2017 ×64 (Solidworks, Waltham, MA) and converted to .TAP files with Sprutcam (Sprutcam, Naberezhnye Chelny, Russia) or to .simpl files with Fusion 360 (Autodesk, San Rafael, CA). Devices were milled from 2 mm thick polystyrene (PS) sheets (Goodfellow USA, Coraopolis, PA). After milling, the device was washed thoroughly with dish soap and water, followed by 70% ethanol and DI water then air dried. A piece of 1.2 mm PS plastic (Goodfellow USA, Coraopolis, PA) was cut (65×50 mm square), cleaned, and solvent bonded to the bottom surface of the device according to the protocol adapted from Young et al. (60). Briefly, a hot plate (Torrey Pines Scientific, HP60) was pre-heated to 65 °C; the bottom of the milled device was mated to the top surface of 1.2 mm PS square; small amounts of acetonitrile (ACN; A998-4, Fisher Chemicals) was pipetted gently through the hollow regions of the device until the liquid filled the entire mated area between the PS square and the device. Weight (2 kg) was placed on the top for about 30 s to facilitate the reaction between solvent and the polymer interface. The bonded device was sonicated with 70% ethanol at 68 °C for 15 min and rinsed with DI water and air dried. The device was plasma treated for 5 min at 0.25 mbar and 70 W in a Zepto LC PC Plasma Treater (Diener Electronic GmbH, Ebhausen, Germany) using oxygen, followed by 10 min of UV sterilization. An engineering drawing of the device with dimensions is included in SI Fig. 1. Original design files are included in the ESI.

### Isolation and cell culture of HPTEC and HKMEC

This work was approved by the University of Washington Institutional Review Board (IRB447773EA). Human HPTECs and HKMECs were isolated from surgically dissected kidney tissues from fetal donors, as we reported previously (33). Kidneys were minced in serum-free EBM2 endothelial growth medium (Lonza) with 0.2 mg/mL Liberase and 100 U/mL DNase (Roche). The minced kidneys were incubated for 30 min at 37 °C in a shaking water bath. The tissue homogenate was filtered through a 40 *μ*m cell strainer. For HKMEC enrichment, EpCAM-positive cells were depleted first from the cell suspension using EpCAM microbeads (Miltenyi Biotec). EpCAM-negative cells were cultured on gelatin-coated T-75 flasks in EBM2 media supplemented with antibiotic/antimycotic (Invitrogen), 10% FBS (Invitrogen), 10 *μ*g/mL ECGS (Cell Biologics), 50 ug/mL heparin (Sigma), and 40 ng/mL VEGF (R&D Systems). Cells were cultured at 5% CO_2_ and 37 °C until confluent, and HKMECs were sorted as CD144-positive population on BD FACSAria II at the UW SLU Flow Cytometry Facility. After sorting, HKMECs were cultured in EBM2 media supplemented with antibiotic/antimycotic (Invitrogen), 10% FBS (Invitrogen), 10 *μ*g/mL ECGS (Cell Biologics), 50 *μ*g/mL heparin (Sigma), and 20 ng/mL VEGF (R&D Systems) at 5% CO_2_. For HPTEC enrichment, EpCAM-positive cells were collected using EpCAM microbeads from the cell suspension. HPTECs were then cultured in T-75 flasks in DMEM/F12 (Corning) supplemented with antibiotic/antimycotic (Invitrogen), ITS-X (Invitrogen), and 50 nM hydrocortisone (Sigma-Aldrich) at 5% CO_2_ and 37 °C. Cells between passage 1-5 were used for coculture experiments.

HKMECs and HPTECs were plated on the center (17 *μ*L) and side chamber (37 *μ*L) of the coculture device at a density of 1×10^5^ cells/cm^2^ and 4.35×10^4^ cells/cm^2^ with their own media, respectively. 24 hours after initial cell plating, the two chambers were connected using 100 *μ*L HKMEC media +/− VEGF (20 mg/mL), which was replaced every 24 h. Throughout culturing, the devices were kept in a primary container (a 86×128 mm OmniTray, or a 100×150 mm Petri Dish (Thermo Scientific)). Sacrificial PBS droplets (total volume of 1 mL) were pipetted around the devices to mitigate evaporation. The primary container was placed in the center of a 245×245 mm BioAssay Dish (Corning) with 100 mL PBS. The peripheral of the primary container was wrapped tightly with moisten Kim Wipe. The secondary container was incubated at 5% CO_2_ and 37°C.

For HKMECs and HPTECs conditioned media culture experiment, HPTECs were plated on the side chambers of the device and recovered overnight. On the next day, HPTECs were fed with 100 *μ*L HKMEC media supplemented with 20 mg/mL VEGF. HKMEC were plated on the center chambers of separate wells. After overnight recovery, HPTEC conditioned media was collected and filtered to remove any HPTEC debris. 100 *μ*L of conditioned media was added to the wells containing HKMECs. The conditioned media was collected and changed every 24 h.

### Cell culture of HUVEC and hPSC-EC

HUVECs (Lonza) were cultured in EBM (Lonza) supplemented with EGM Endothelial Cell Growth Medium SingleQuots Kit (Lonza) and used at passage 3-6. The coculture procedures for HUVEC-HPTEC were the same as HKMEC-HPTEC coculture described above using HUVEC media. Human pluripotent stem cells (hPSC) (WTC line, a gift from Dr. Murry, University of Washington) was first differentiated to posterior-like ECs according to the protocol adapted from Palpant et al. (41) and Redd et al. (44) with low Activin A and high BMP4 for 10 days. Then these cells were trypsinized and plated into co-culture device (at center well) to allow for attachment and growth into confluency. HPTECs were plated into the outer well of the device at day 11, and co-cultured with hPSC-EC for additional two days in hPSC-EC media (41). At day 13, five devices were fixed with PFA for immunostaining, and additional 5 devices were lysed for RNA collection.

### Cell staining and imaging

The ECs in the study were washed with PBS, blocked with PBS containing 2% BSA for 1 h, and incubated with 0.1% Triton X-100 for 5 min. The cells were washed and incubated at 1:50 dilution over night at 4°C with primary antibodies: mouse anti PV-1, rabbit anti VECad, and sheep anti vWF (Abcam). The samples were washed and incubated at 1:200 dilution for 1 hour at room temperature with secondary antibodies: goat anti mouse 488, goat anti rabbit 568, and donkey anti sheep 568 goat anti-rat secondary antibody (Thermo Fisher). The secondary antibodies were removed, a 1:1000 dilution of DAPI (Invitrogen) was added and incubated for 5 min. The samples were washed twice, mounted with PBS, and covered with a glass slip to prevent sample dehydration during imaging. The protected device was fitted directly into the mounting frame for fluorescence microscopy.

### RNA isolation, RT-PCR, and qPCR analysis for ECs

RNA was collected and purified using the RNAeasy Mini Kit (Qiagen), and residual DNA was removed by on-column DNase digestion (Qiagen). RNA quantity and quality were measured with NanoDrop ND 1000 Spectrophotometer. RT-PCR was performed using Biometra T-Personal Thermal Cycler with iScript Reverse Transcription Supermix (Bio-Rad). qPCR was performed using the Real-time PCR System (Applied Biosystems) with Fast SYBR Green Master Mix (Applied Biosystems) and primers (RealTimePrimers). The gene expression was quantified according to MIQE guidelines, and the expression of each gene was determined relative to reference gene GAPDH (10).

### Microscopy and image processing

ECs were imaged using a Nikon High Resolution Widefield microscope with a high-resolution CCD camera. Brightness/contrast adjustments were made uniformly across all images in Fiji (ImageJ). To calculate the cell size for HKMECs monoculture without VEGF condition, a random cell region on 10x
image was chosen and cells were counted, and average cell size was determined by region area/DAPI count. For other culture conditions where HKMECs were confluent, the average cell size was determined by field of view/total DAPI count.

### Statistical analysis

Quantitative data was graphed and analyzed using GraphPad Prism 7 software. In Fig. 2C, each plotted point represents the average of approximately 3.5×10^4^ cells – 8.0×10^4^ cells analyzed within a device. HKMEC cell size was compared using a one-way an ANOVA test followed by Tukey's multiple comparisons. In Figs. 3 and 4, statistical tests were run to compare the expression in coculture vs. monoculture (all VEGF-free) as follows: Due to the distribution of data from multiple human donors, the coculture expression ratios for the three donors were log transformed to better fulfill the assumptions of the *t*-test. The log transformed coculture data were then compared to 0 using a two-tailed one-sample *t*-test using “QuickCalcs one-sample *t*-test” on the GraphPad website. (Note that the coculture data had previously been normalized to monoculture data, which was set to 1 as described in the figure captions; log(1)=0 hence we compared to 0 using the one-sample *t*-test.)

## Acknowledgements

We thank John Day and Jing Lee for device milling support, and Sam Rayner for imaging support.

## Author contribution

Y.Z., A.B.T., and J.H. conceived and supervised the biological aspects of the project. A.B.T., E.B., and T.Z. conceived and designed the microfluidic coculture device. Y.Z., A.B.T., T.Z., D.L. designed the experiments. T.Z. and D.L. performed the experiments with assistance of R.J.N. and J.X. Results were analyzed by T.Z., D.L., Y.Z., J.H., and A.B.T., and T.Z., Y.Z., and A.B.T. wrote the manuscript. All authors approved the study.

