## Supplemental Information for "Open microfluidic coculture reveals paracrine signaling from human kidney epithelial cells promotes kidney specificity of endothelial cells" (from another version of this preprint: version 2, 2020-02-19)

***Supplementary Material***

Tianzi Zhang^1^, Daniel Lih^2^, Ryan J. Nagao^2^, Jun Xue^2^, Erwin Berthier^1^, Jonathan Himmelfarb^*3,4^, Ying Zheng^*2,4,5^, and Ashleigh B. Theberge^*1,4,6^

Departments of ^1^Chemistry, ^2^Bioengineering, ^3^Medicine, ^4^Kidney Research Institute, ^5^Institute for Stem Cell and Regenerative Medicine, and ^6^Urology, University of Washington, Seattle, 98105, USA

**Spontaneous capillary flow (SCF) in the device side culture chamber**

In an open system fluid flow can occur without the need for external pressure sources, which is described as spontaneous capillary flow (SCF). (22, 24, 33) SCF occurs when the following inequality is fulfilled:

$$\frac{P_{f}}{P_{w}}<\cos\left( \theta\right)$$

where the wetted perimeter (*P_w_*) comprises all the solid wetted surfaces in the channel cross-section, the free perimeter (*P_f_*) comprises all the open air-liquid interfaces in the cross-section, and 𝜃 represents the contact angle of the liquid on the channel material. In the microfluidic coculture device described in this manuscript, the side chamber cross sectional geometry (Figure S1C) fulfills the conditions for spontaneous capillary flow as shown below. The contact angle of cell culture media on device surface (plasma treated polystyrene) is 17.9°±3.0°. (24)

$\cos\left( 17.9^{\circ} \right)\approx0.95$,

$$\frac{P_{f}}{P_{w}}=\frac{1.6 mm}{(2.0+1.5+1.4) mm}\approx0.33$$

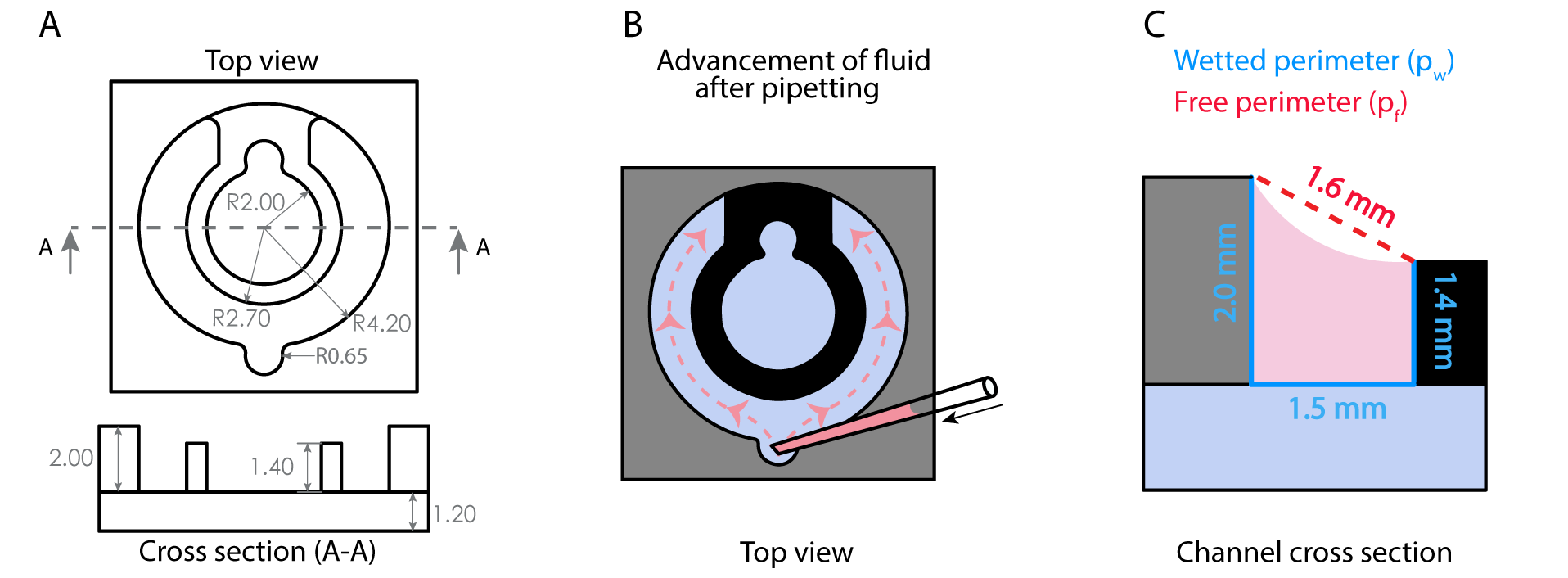

Figure S1. Device dimensions and spontaneous capillary flow (SCF) during the microscale device loading. (A) Top view and cross section illustrating the dimensions of the coculture platform. Unit: mm. Computer aided design (CAD) file is included in the ESI. (B) Top view showing the advancement of fluid in the device side channel. (C) Channel cross section geometry showing the wetted perimeter (*p_w_*) and free perimeter (*p_f_*), which fulfills the condition for SCF.

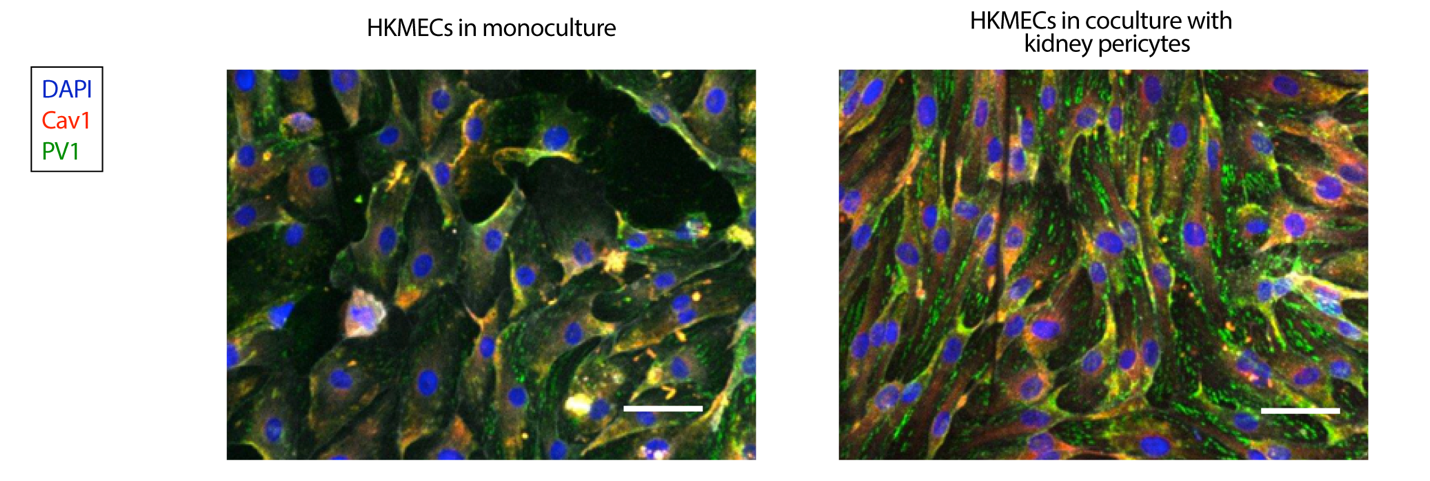

Figure S2. Incorporation of kidney pericytes increased HKMEC density in coculture. Blue: nuclear stain (DAPI). Red: Caveolin- 1 (Cav1); Green: plasmalemma vesicle associated protein (PV1). Scale bars: 50 μm.

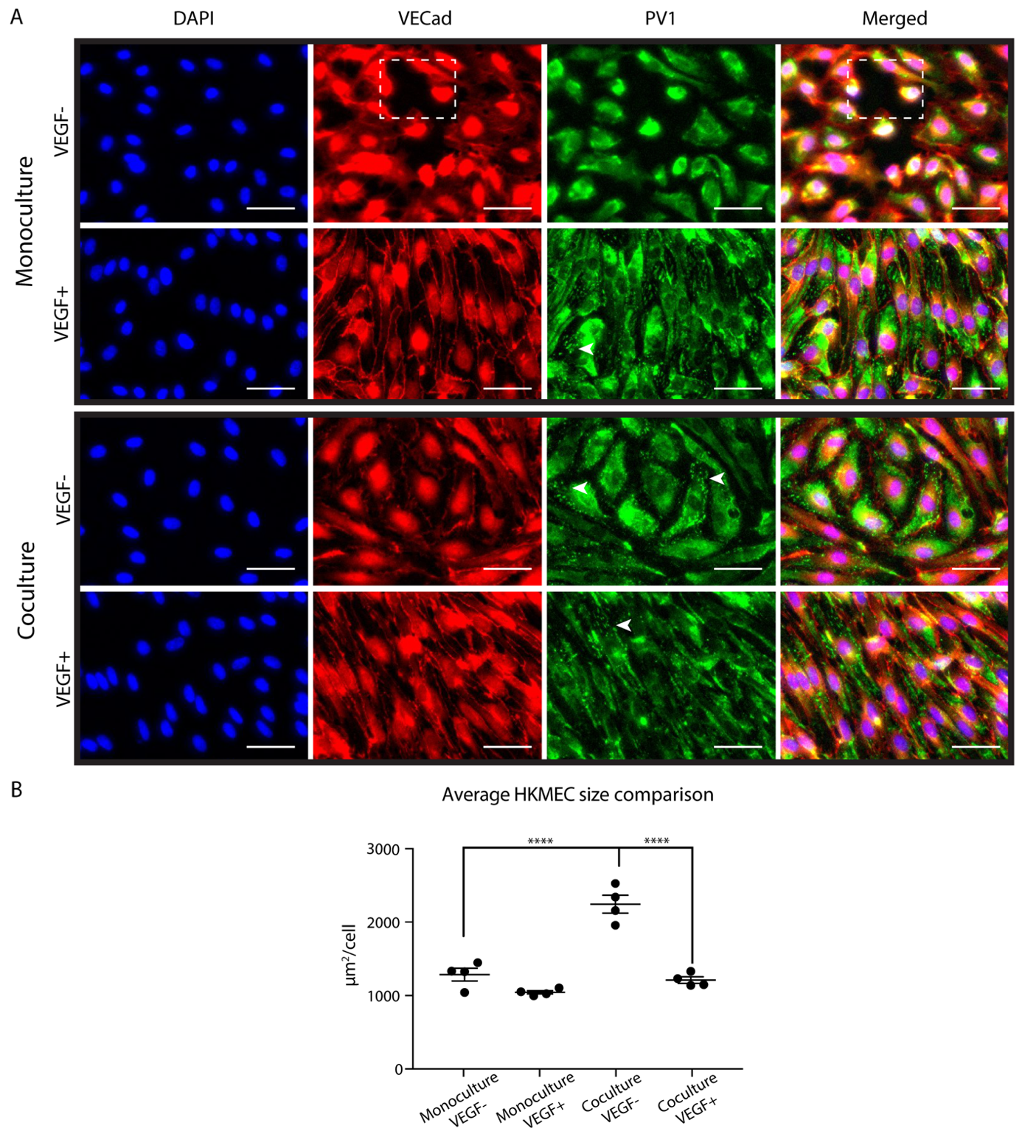

Figure S3. Coculture with HPTECs preserves HKMECs morphology in coculture. (A) Immunofluorescence images of HKMECs under differential culture conditions. Blue: nuclear stain (DAPI). Red: vascular endothelial cadherin (VECad); dotted box indicates a region without cells. Green: plasmalemma vesicle associated protein (PV1); arrows indicate example regions with clear PV1 structures (punctate staining). Scale bars: 50 μm. Images shown are from donor B and are representative of experiments with cells from 3 different human donors each with 5 device replicates per experiment. (B) Quantification of HKMEC size under differential culture conditions. Error bars represent the SEM of five device replicates for a single human donor (donor A) in a single experiment. Data sets were analyzed using a one-way ANOVA test; *p*-values are indicated for Tukey’s multiple comparisons tests between “Coculture, VEGF-” and “Monoculture, VEGF-” conditions; and “Coculture, VEGF-” and “Coculture, VEGF+” conditions. (*****p* ≤ 0.0001)

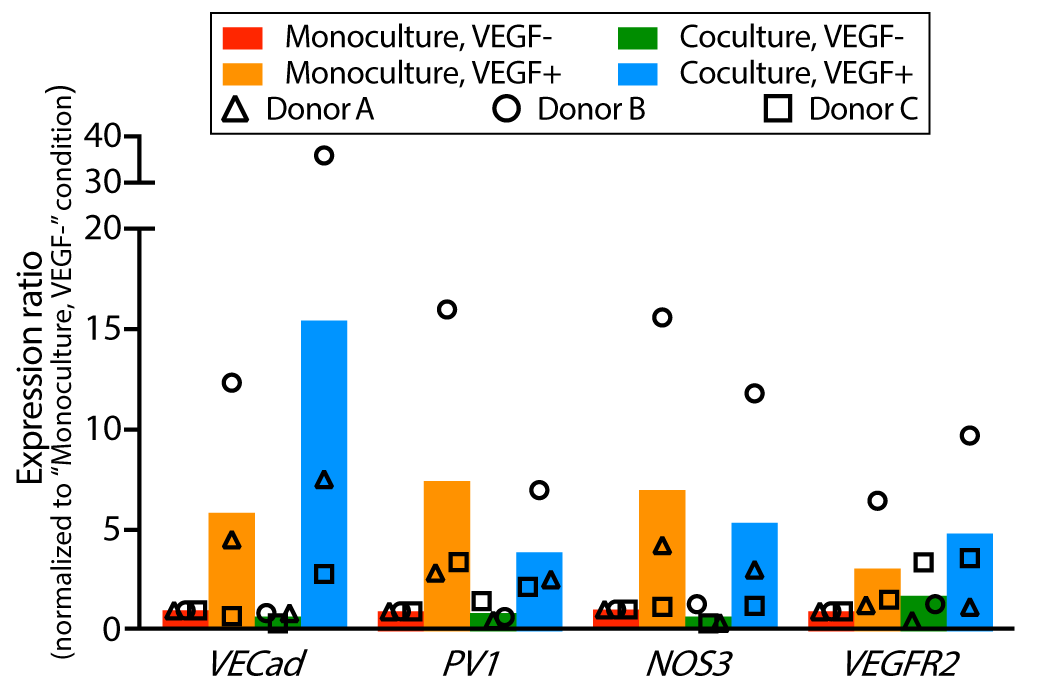

Figure S4. RT-qPCR data showing downregulation of selected HKMEC genes in VEGF-free media (red and green bars), suggesting an important role of VEGF in endothelial cell growth and development. Each plotted point represents data from an independent human donor (pooled from 5 replicate microculture devices per donor, average of 3 RT-qPCR technical replicates), with each normalized to the “Monoculture, VEGF-” condition (which is set to 1). Each colored bar represents the average ratio of the three donors.

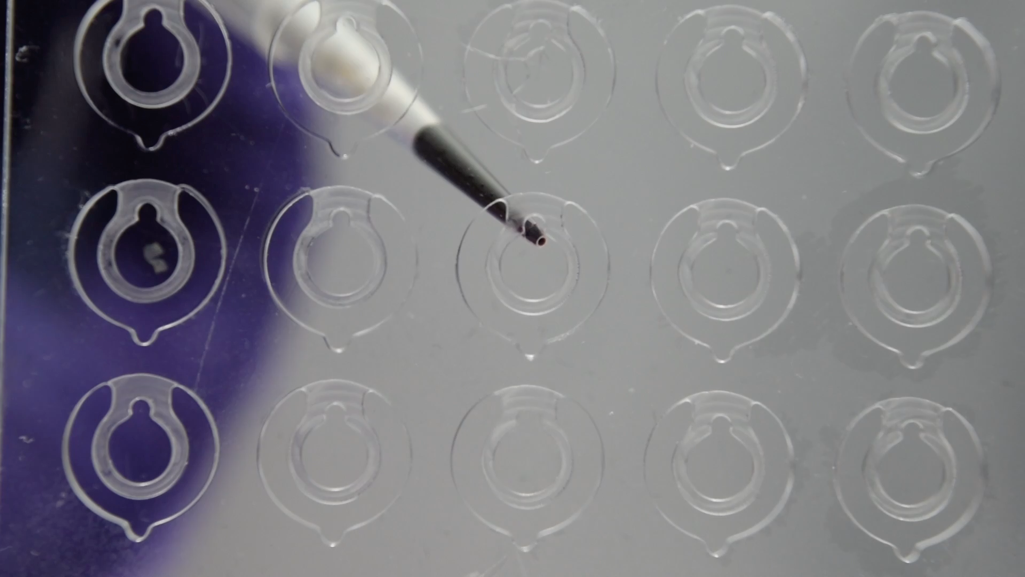

Video S1. Coculture device loading procedure. Purple dye solution (17 μL) and green dye solution (37 μL) were selectively pipetted to the center and side chamber of a well on the coculture device. Water (46 μL) was then loaded to connect the two chambers (The total loading capacity of the well is 100 μL).
